# Dynamic subcellular localization of sodium-bicarbonate cotransporter NBCn1/SLC4A7 to plasma membrane, centrosomes, spindle, and primary cilia

**DOI:** 10.1101/2022.10.05.510992

**Authors:** Marc Severin, Emma Lind Pedersen, Magnus Thane Borre, Ida Axholm, Frederik Bendix Christiansen, Muthulakshmi Ponniah, Dominika Czaplinska, Tanja Larsen, Luis Angel Pardo, Stine Falsig Pedersen

## Abstract

Finely tuned regulation of transport protein localization is vital for epithelial function. Sodium-bicarbonate co-transporter NBCn1 (SLC4A7) is a key contributor to epithelial pH homeostasis, yet the regulation of its subcellular localization is not understood. Here, we show that a predicted N-terminal β-sheet and short C-terminal α-helical motif are essential for NBCn1 plasma membrane localization in epithelial cells. This localization was abolished by cell-cell contact disruption, and co-immunoprecipitation (co-IP) and proximity ligation (PLA) revealed NBCn1 interaction with E-cadherin and DLG1, linking the transporter to adherens junctions and the Scribble complex. NBCn1 also interacted with RhoA and localized to lamellipodia and filopodia in migrating cells. Finally, analysis of localization of native and GFP-tagged NBCn1, subcellular fractionation, co-IP of NBCn1 with Arl13B and CEP164, and PLA of NBCn1 and tubulin in mitotic spindles led to the surprising conclusion that NBCn1 additionally localizes to the centrosome and primary cilium in non-dividing, polarized epithelial cells, and to spindle, centrosome and midbodies during mitosis. We propose that NBCn1 traffics between lateral junctions, leading edge, and cell division machinery in Rab11 endosomes, adding new insight to the role of NBCn1 in cell cycle progression.

**Summary statement:** We unravel molecular determinants of plasma membrane localization of the Na^+^,HCO_3_^−^ cotransporter NBCn1 and discover that NBCn1 also localizes to centrosomes, spindle, midbody and primary cilia, likely cycling between these compartments.

## INTRODUCTION

Finely tuned regulation of the subcellular localization of ion transport proteins is vital for tissue function, a notion substantiated by the fact that mutations causing ion transporter mislocalization underlie many human diseases (Almomani et al., 2011, Denning et al., 1992, Wilson et al., 1991). Sodium-bicarbonate co-transporter NBCn1 (SLC4A7), a member of the SLC4 family of bicarbonate transporters, is a major net acid extruder with essential physiological roles in many human organs. NBCn1 is widely present in both epithelial and non-epithelial cells (Boedtkjer et al., 2012, Parker and Boron, 2013) and its dysregulation is implicated in major diseases such as hypertension and cancer (Ng et al., 2017, Gorbatenko et al., 2014a, Boedtkjer, 2019). Consistent with the delayed growth rate of *in vivo* tumors and tumor cells with reduced NBCn1 expression (Andersen et al., 2018, Lee et al., 2016), knockdown of NBCn1 in MCF-7 breast cancer cells substantially delayed cell cycle progression in synchronized cells. This delay was most pronounced during G2/M transition, where NBCn1 expression also peaked, suggesting a particularly important role for the transporter upon entry into mitosis (Flinck et al., 2018).

Based on homology to SLC4 family transporters with available cryo-EM structures (Arakawa et al., 2015, Huynh et al., 2018, Wang et al., 2021), NBCn1 is expected to comprise ~14 transmembrane helices, a long N-terminal and a short C-terminal cytosolic domain, and to form homodimers in the membrane. The ~600 amino acid N-terminal cytosolic domain comprises a proximal part predicted to be largely unfolded, followed by two core interacting regions linked by a variable loop (Parker and Boron, 2013). In epithelial tissues, NBCn1 primarily localizes to the basolateral membrane (Parker and Boron, 2013). However, in the choroid plexus, NBCn1 is apically localized (Damkier et al., 2007), and in subconfluent, migratory cells, NBCn1 localizes to lamellipodial structures (Olesen et al., 2018), suggesting that cell type and polarity impacts transporter localization.

The C-terminal cytosolic tail of NBCn1 is essential for its membrane localization (Loiselle et al., 2003, Olesen et al., 2018) and links the transporter to several membrane proteins via a PDZ binding motif (Lee et al., 2014). Through a pulldown with NBCn1’s C-terminal tail followed by mass spectrometry (MS) analysis we recently identified several putative NBCn1 binding proteins involved in membrane retention, trafficking, and degradation of the transporter (Olesen et al., 2018). Of these, the scaffold protein Receptor for activated C-kinase (RACK1) interacted with the most proximal part of the NBCn1 C-terminus and was important for trafficking of NBCn1 to the plasma membrane. We also found that NBCn1 is constitutively endocytosed from the basolateral membrane but only very slowly degraded, indicative of recycling (Olesen et al., 2018). However, the specific motifs involved in controlling NBCn1 membrane localization, and the signaling events triggering changes in its membrane localization are unknown. Furthermore, in addition to the expected trafficking- and plasma membrane-localized NBCn1 binding candidates, our MS analysis identified proteins associated with lamellipodia/filopodia formation, cell-cell junction and polarity complex proteins, as well as some fully unexpected putative partners of high confidence, localised to centrosomes and primary cilia.

To understand these surprising findings, we conducted a detailed investigation of the subcellular localization of NBCn1 and the molecular motifs and mechanisms controlling it. We show that both the N- and C-terminal cytosolic regions of NBCn1 are essential for its plasma membrane localization, and we identify the specific NBCn1 regions and motifs involved. We demonstrate that NBCn1 interacts tightly with cell-cell adhesion proteins, epithelial polarity proteins and proteins essential to lamellipodia-/filopodia formation, likely underlying the potent regulation of its membrane localization by cell confluence. Finally, we demonstrate, using multiple different lines of evidence, that under specific conditions, NBCn1 localizes to primary cilia, centrosomes and mitotic spindle of epithelial cells. This work places NBCn1 among the growing number of ion transport proteins with highly non-canonical localizations and functions and provides a new framework for understanding the pivotal role of NBCn1 in cell function.

## RESULTS

### A short proximal C-terminal α-helical motif is essential for NBCn1 plasma membrane localization

Because we previously demonstrated that RACK1 interacted with the most proximal part of the C-terminal NBCn1 tail (C-tail) (Olesen et al., 2018), we suspected that key motifs important for NBCn1 trafficking would localize here, and we focused our efforts on this region. Informed by protein fold predictions (Fig. 1A), Alphafold structure (Fig. 1B), and sequence conservation between species (Suppl. Fig. 1A) and across SLC4A family members (Suppl. Fig. 1B), we designed and generated a series of GFP-NBCn1 C-terminal truncations and deletion constructs of the rat NBCn1-D isoform, which we transiently expressed in MCF-7 human breast cancer cells. For clarity, Fig. 1 includes a small part of the last α-helix of the NBCn1 transmembrane domain (TMD). The proximal C-terminal cytosolic domain contains two predicted α-helical stretches (marked I-II), of which the first is very short (Fig. 1A, B). Interestingly, helix I contains the highly conserved motif RELSWLD (Suppl. Fig. 1A-B), which is found in several otherwise unrelated proteins, suggesting it may serve a regulatory function (see Discussion).

**Figure 1.**
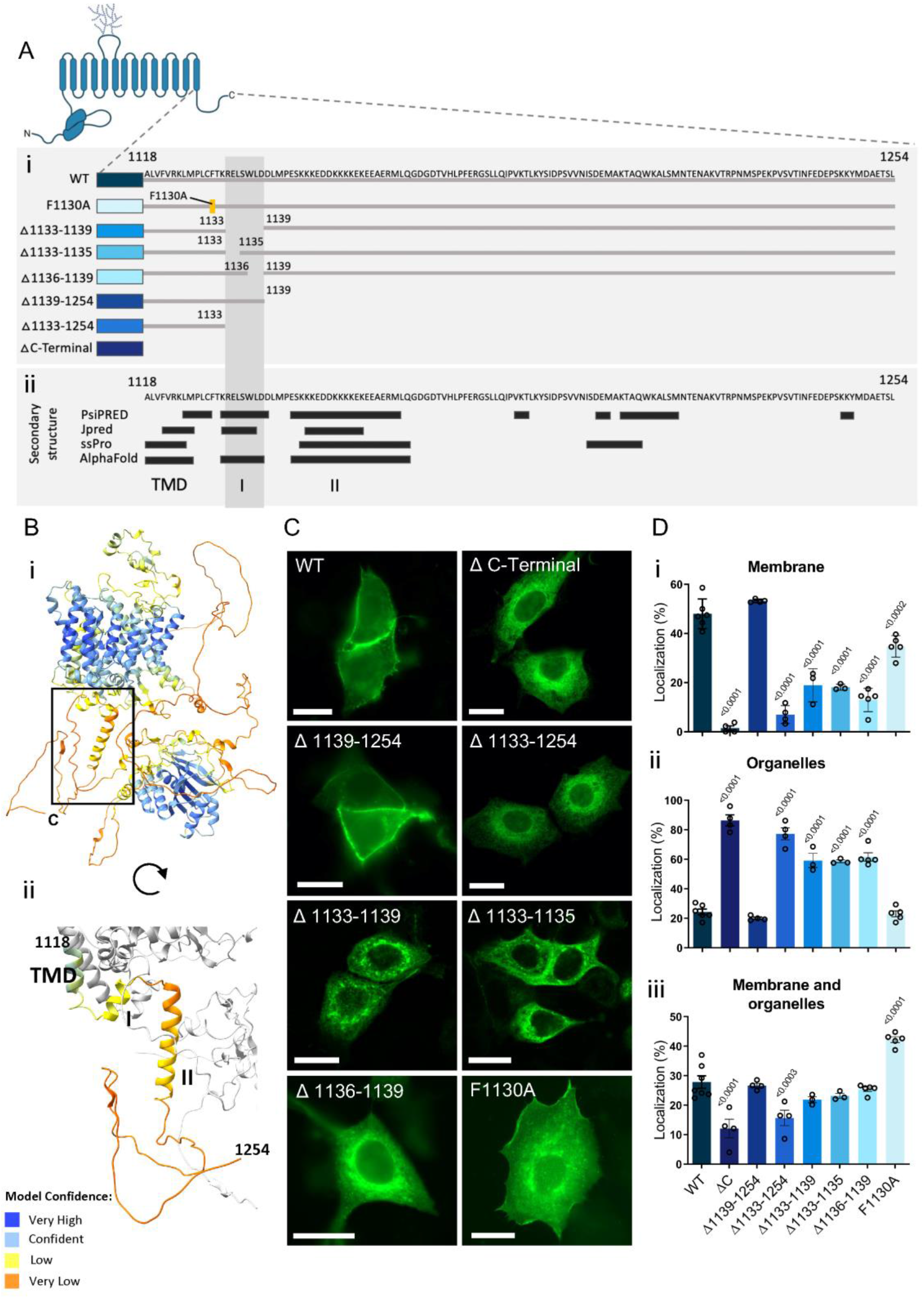
C-terminal determinants of NBCn1 plasma membrane localization. **(A)** Overview of the C-terminal cytosolic tail of NBCn1 and the constructs generated. The NBCn1 sequence is *R. norvegicus* NBCn1-D (NCBI ref: NM_058211.2), highly similar to human NBCn1 (Suppl. Fig. 1A). Secondary structure predictions were made using PSIPred (Buchan and Jones, 2019), JPred4 (Drozdetskiy et al., 2015) and SSPro (Urban et al., 2022). The three helical regions (black bars) correspond to part of the last helix of the transmembrane domain (TMD), and the proximal (I) and distal α-helix (II), the grey bar marks the RELSWLD motif (elm.eu.org, (Kumar et al., 2021)). (B) Predicted structure of the human NBCn1 C-terminal (Uniprot ID: Q9R1N3) from Alphafold 2 (Jumper et al., 2021). Zoom in ii shows the localization of helices I and II. **(C)** MCF-7 cells were transiently transfected with GFP-NBCn1 constructs expressing full length or variant NBCn1 as indicated and imaged with fluorescence microscopy. Scale bars: 20 μm. **(D)** NBCn1 plasma membrane localization was analyzed as described in Materials and Methods, and data are shown as localization to plasma membrane (i), organelles (ii), or both (iii). n=3-6. Statistical analysis: one-way ANOVA with Dunnett’s multiple comparisons test.

Consistent with earlier work (Olesen et al., 2018, Loiselle et al., 2003), deletion of the full C-terminal tail abolished NBCn1 plasma membrane localization (Fig. 1C-D). As we predicted from our RACK1 interaction analysis (Olesen et al., 2018), truncation of the entire C-terminal distal to α-helix I (i.e. Δ1139-1254) had no effect on NBCn1 plasma membrane expression (Fig. 1C-D). In marked contrast, a truncation including just the six additional amino acids of helix I (Δ1133-1254) essentially abolished plasma membrane localization. Strikingly, deletion of helix I alone (1133-1139) reduced plasma membrane localization by nearly two-thirds, and deletion of either residues 1133-35 or 1136-39 of the α-helix had a similar effect. A phenylalanine residue at 1130 that is part of a predicted cholesterol-binding CARC motif (Fantini and Barrantes, 2013) (^1124^<u>K</u>LMDLC<u>F</u>TKRE<u>L</u>^1135^), did not appear to play a role, as a point mutation to alanine (F1130A) had no discernible effect on NBCn1 localization (Fig. 1C-D).

Integral membrane proteins destined for the basolateral membrane can take several possible routes from the trans-Golgi network (TGN) to the plasma membrane, either directly or via recycling endosomes (REs). This often involves the epithelial-specific Adaptor Protein, AP-1B, which in polarized MDCK cells localizes to REs (Cancino et al., 2007, Farr et al., 2009). On the other hand, some basolateral proteins including E-cadherin (Miranda et al., 2001) and the Na^+^,K^+^-ATPase (Farr et al., 2009) do not depend on AP-1B for basolateral targeting. We noted that multiple components of the AP1 complex (AP-1μ1, −μ2, −γ, −σ, and −β) were pulled down with NBCn1 in our MS data (Suppl. Fig. 2A). Indeed, interaction of NBCn1 with the AP-1 complex was confirmed by native co-immunoprecipitation (co-IP) of NBCn1 with γ-adaptin in both confluent (100%) and non-confluent (70%) MDCK II cells (Suppl. Fig. 2B), consistent with the interpretation that NBCn1 trafficking to the basolateral membrane may at least in part involve an AP-1B- and RE-dependent pathway.

These results show that NBCn1 plasma membrane localization is fully dependent on six amino acids in a small helical motif in the proximal part of the C-terminal cytosolic tail of the transporter and may involve an AP-1B – RE pathway.

### An N-terminal predicted β-sheet is important for NBCn1 plasma membrane localization

We next performed a similar analysis of the N-terminal cytosolic domain, focusing on the first 148 residues because of the conspicuous motifs and predicted structure in this region (Fig. 2A, Suppl. Fig. 1C). Alphafold predicts a short α-helical domain at residues 95-104 and a β-sheet at 117-146, immediately before the core domain (Fig. 2A, B). Two lysines in this region, K135 and K142, caught our attention because they are contained in predicted sumoylation motifs (Flotho and Melchior, 2013) and in a motif corresponding to a suggested regulatory, Zn^2+^ binding motif in SLC4A8 (NDCBE) (Alvadia et al., 2017).

**Figure 2.**
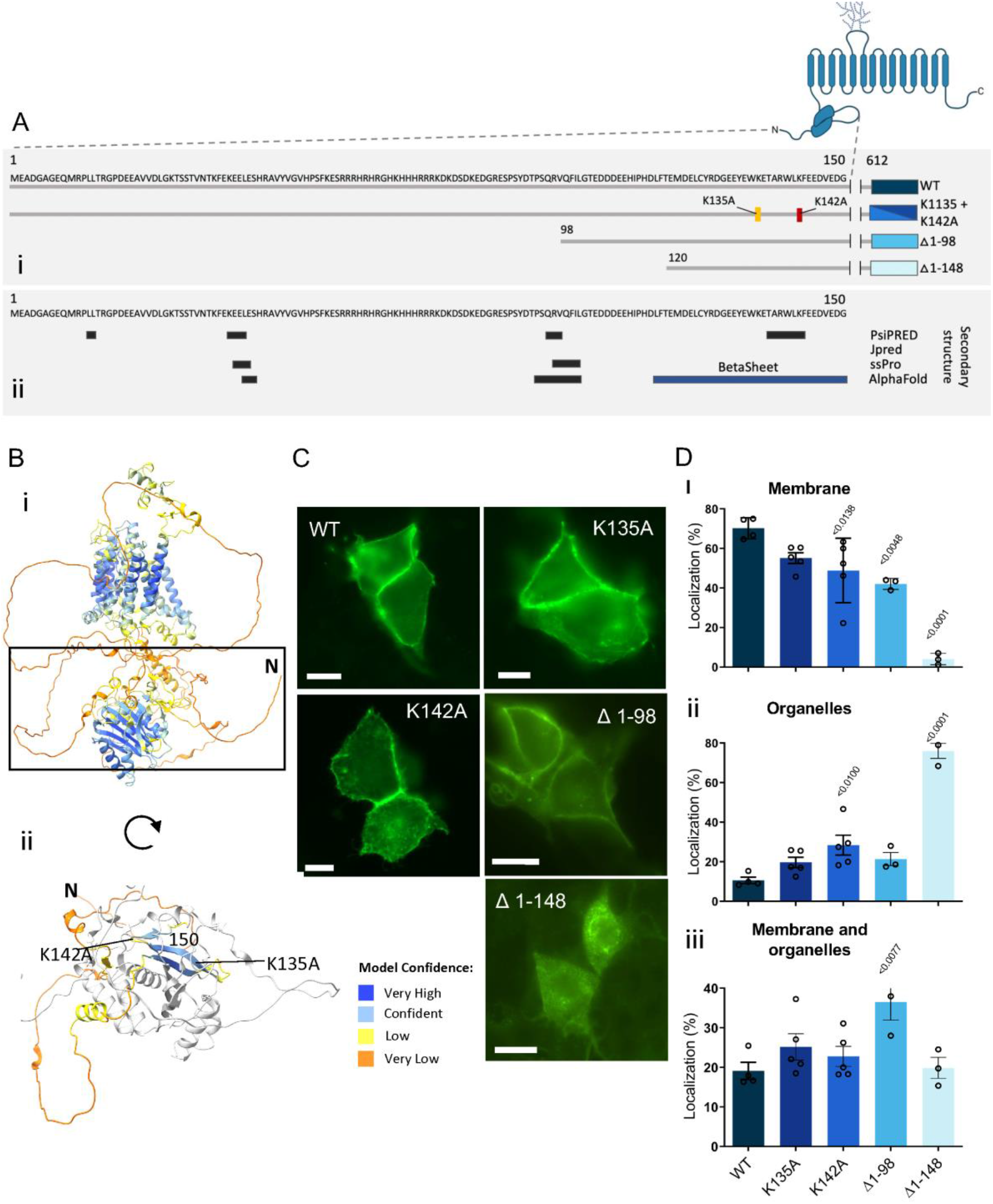
N-terminal determinants of NBCn1 plasma membrane localization. **(A)** Structure of the N-terminal domain of human NBCn1 (Uniprot ID: Q9R1N3), showing the predicted β-sheet based on predictors as in Fig. 1 **(B)** Predicted structure of NBCn1’s C-terminal from Alphafold 2. **(C)** MCF-7 cells were transiently transfected with NBCn1-GFP constructs expressing either full length NBCn1 or gene with deletion or point mutations and imaged with fluorescence microscopy. Scale bars: 20 μm. **(D)** Plasma membrane localization was quantified as membrane localization (i), organelle localization (ii), or a combination of the two (iii) (n=3). Statistical analysis: one-way ANOVA with Dunnett’s multiple comparisons test.

We therefore generated four full-length NBCn1 variants: K135A, K142A, Δ1-98, and Δ1-148 and examined their membrane localization after expression in MCF-7 cells (Fig. 2C-D). The K135A mutation had no effect, K142A mutation or truncation of the first 98 residues (Δ1-98) both reduced plasma membrane localization by about one-third, while additionally removing the predicted β-sheet (Δ1-148) almost completely abrogated NBCn1 plasma membrane expression (Fig. 2C-D). This pattern was stable over a time course of 96 h, and hence did not appear to be simply a matter of slower kinetics of transport (Suppl. Fig. 2C). NBCn1 variants with impaired plasma membrane localization localized in part to an intracellular compartment staining for Golgin-97 and giantin, consistent with accumulation in both the Trans-Golgi Network (TGN) and cis-medial Golgi (Suppl. Fig. 3D).

These findings show that NBCn1 membrane localization is dependent not only on the short C-terminal motif, but also on the proximal part of the N-terminal, including a predicted β-sheet.

### Plasma membrane localization and expression of NBCn1 are induced by cell confluence

Basolateral delivery of vesicles containing integral membrane proteins often involves their tethering to the exocyst complex at apical-lateral junctions before distribution throughout the basolateral plasma membrane (Rodriguez-Boulan et al., 2005). Such a mechanism would imply that NBCn1 targeting to the membrane is dependent on some degree of pre-existing polarity. To examine this, MDCK II cells cultured for 24 h to either 20% or 60% confluence or for a week at 100% confluence were subjected to immunocytochemistry (ICC) analysis for NBCn1, with E-cadherin as a plasma membrane marker (Fig. 3A-C). Remarkably, the subcellular localization of NBCn1 changed completely with increasing confluence: in non-confluent conditions, i.e. in absence of epithelial polarization, the transporter was distributed throughout the cells, including the nucleus/nuclear membrane (Fig. 3A-B). In the confluent, polarized cells, NBCn1 was almost exclusively found at the plasma membrane (Fig. 3C). These results were validated by line scan quantification, with examples from each condition shown in Fig. 3D-F and summary data in Fig. 3G. As seen, relative NBCn1 membrane localization increased with increasing cell confluence, and was approximately doubled in fully confluent cells compared to 20% confluent cells (Fig. 3G).

**Figure 3.**
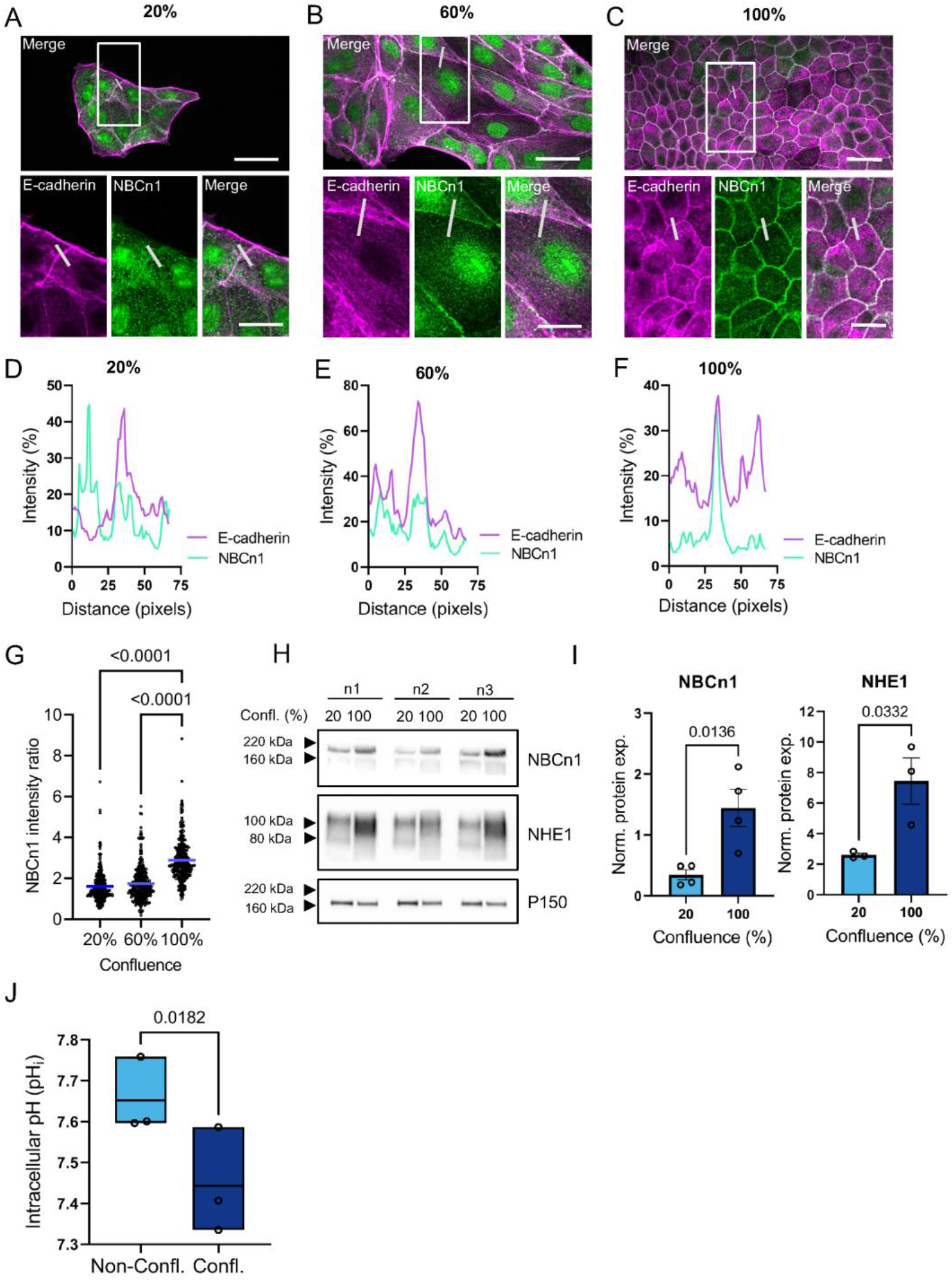
NBCn1 localization depends on cell confluence. (MDCK II cells were cultured for 24 h to 20% **(A)** or 60% **(B)** confluence or for a week at full confluence **(C)** and fixed. Scale bars:.20 μm, zooms: 10 μm. Cells were stained for NBCn1 and E-cadherin as a plasma membrane marker and relative NBCn1 plasma membrane localization was determined through line scan analysis **(D-F)**, summarized in G (n=3). Statistical analysis: one-way ANOVA with Dunnett’s multiple comparisons test. **(H)** Total NBCn1 and NHE1 protein expression under the same conditions was determined through western blot and quantified, normalized to loading control **(I)** (n=3-4). Statistical analysis: Unpaired, two-tailed Student’s t-test. pH_i_ of confluent and non-confluent MDCK II cells measured with SNARF under epifluorescence microscopy (n=3). Statistical analysis, paired, two-tailed Student’s t-test.

Also total cellular NBCn1 expression exhibited a strong dependence on cell confluence, with an almost 4-fold increase in relative NBCn1 protein expression from 20% to 100% confluent cells (Fig. 3H-I). Interestingly, a similar pattern was seen for another major cellular acid extruder, the Na^+^/H^+^ exchanger NHE1 (SLC9A1) (Fig. 3H-I, Suppl Fig. 3). We therefore asked whether upregulation of acid-base transporters could be driven by a decrease in pH_i_ during polarization and serve to maintain physiological pH_i_ under these conditions. Consistent with such a hypothesis, steady state pH_i_ was significantly decreased in confluent (pH_i_ ~7.43) compared to non-confluent (pH_i_ ~7.64) MDCK II cells (Fig. 3J).

These findings show that expression and plasma membrane localization of NBCn1 and NHE1 are strongly increased as epithelial cells polarize, and that pH_i_ is lower in polarized than in sparse MDCK II cells.

### NBCn1 plasma membrane localization is increased by increases in cellular cAMP level

To further understand the physiological mechanisms that regulate NBCn1 plasma membrane localization, we therefore first asked whether decreasing pH_i_ could drive the observed increase in NBCn1 plasma membrane localization. Lowering extracellular pH (pH_e_) elicits a rapid, lasting decrease in pH_i_ (Michl et al., 2022). Sub-confluent MDCK II cells were exposed to pH_e_ 6.3 growth medium for 2 or 24 h and subjected to ICC analysis for NBCn1 (Fig. 4A). However, this treatment failed to elicit a detectable change in relative NBCn1 plasma membrane expression (Fig. 4B), suggesting that the NBCn1 plasma membrane localization in polarized cells is not driven by pH_i_.

**Figure 4.**
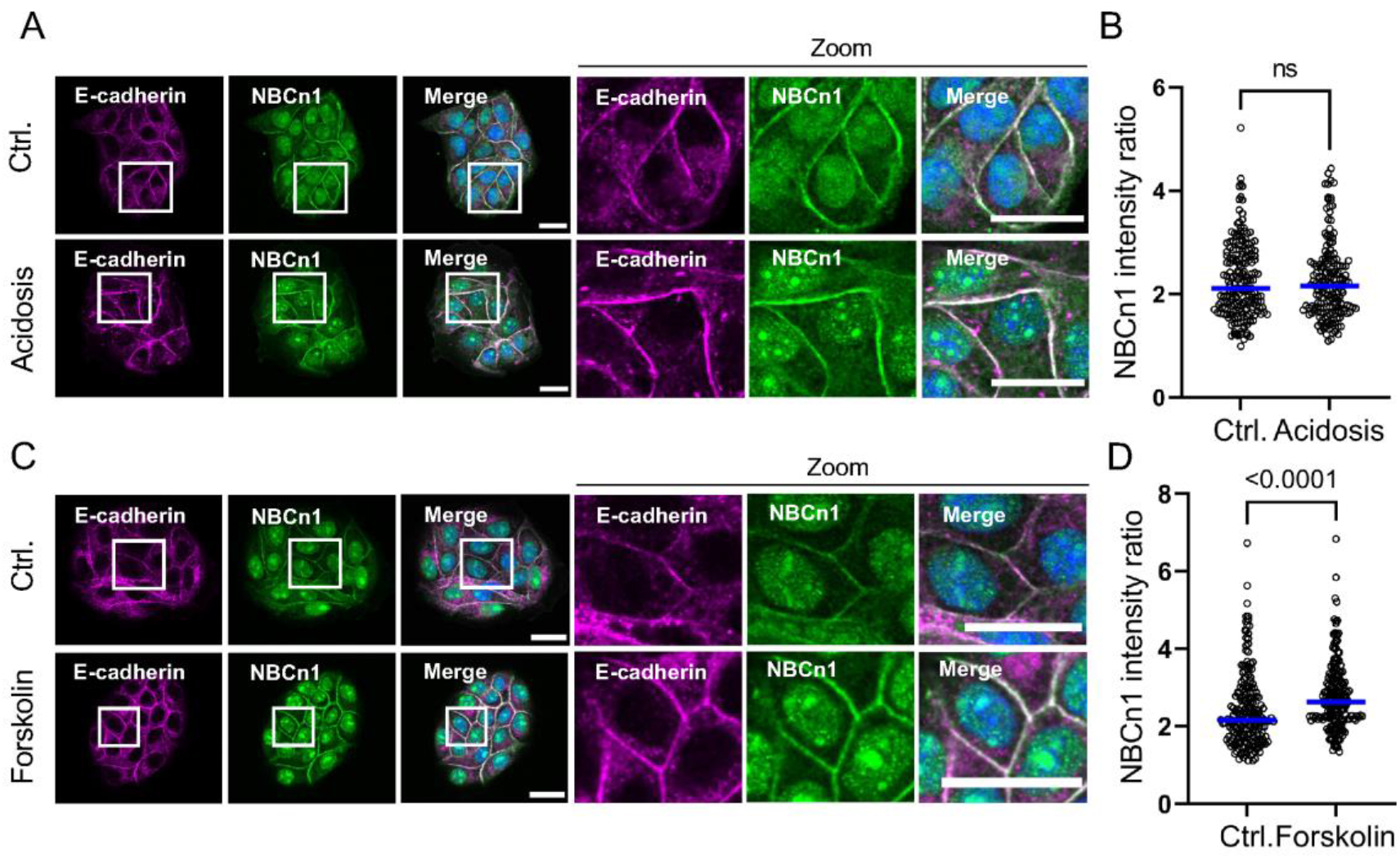
Increases in cAMP, but not short term acidosis, increases NBCn1 membrane localization. **(A)** Sub-confluent MDCK II cells were exposed to 20 μM HCl corresponding to a pH_e_ of ~6.3 in the growth medium for 24 h, fixed and stained for NBCn1 and plasma membrane marker E-cadherin and analysed by ICC. Scale bars: 20 μm. **(B)** Line scan analysis of the intensity ratio of NBCn1 plasma membrane signal between treatments (n=3). Statistical analysis: Mann-Whitney U test **(C)** Sub-confluent MDCK II cells were treated with 20 μM forskolin for 24 h, fixed and stained for NBCn1 and E-cadherin. Scale bars: 20 μm. **(D)** Line scan analysis of intensity ratio between forskolin and control treatment (n=3). Statistical analysis: Mann-Whitney U test.

Epidermal growth factor receptor (EGFR) signaling upregulates NBCn1 expression and acid extrusion capacity in breast cancer cells (Gorbatenko et al., 2014b), and EGFR membrane localization and signaling are confluence dependent (Ranayhossaini et al., 2014, Curto et al., 2007). We therefore asked whether EGFR activation could regulate NBCn1 plasma membrane localization in epithelial cells. Exposure of non-confluent MDCK II cells to 1 μg/mL EGF for 10 or 30 min activated EGFR as detected by increased ERK1/2 phosphorylation (Suppl. Fig. 4A). However, neither 2 nor 24 h of EGF treatment caused a detectable change in NBCn1 plasma membrane localization (Suppl. Fig. 4B-C).

A cAMP-protein kinase A (PKA) signaling axis regulates plasma membrane insertion of many proteins including the related electrogenic Na^+^/HCO_3_^−^ co-transporter NBCe1 (SLC4A4) (Yu et al., 2009, Carranza et al., 1998), and cAMP plays a key role in intracellular HCO_3_^−^ sensing via soluble adenylate cyclase (sAC) (Chen et al., 2000). To study whether cAMP regulates NBCn1 plasma membrane localization, we treated sub-confluent MDCK II cells with the adenylate cyclase activator forskolin. Confirming the activation of PKA, forskolin (20 μM, 10 or 30 min) increased the phosphorylation of the PKA target cAMP response element binding protein (CREB) (Suppl. Fig. 4D). While 2 h of forskolin treatment had no detectable effect (Suppl. Fig. 4E), 24 h forskolin treatment significantly increased relative NBCn1 plasma membrane localization (Fig. 4D).

These data show that in MDCK II cells, NBCn1 plasma membrane localization is increased by a long-term increase in cellular cAMP, but neither by acute extracellular acidification nor by EGFR activation.

### NBCn1 dynamically localizes to lateral domains and interacts with cell polarity and - adhesion proteins

The pronounced increase of NBCn1 plasma membrane localization with increasing epithelial confluence led us to consider whether NBCn1 localization is specifically linked to cell polarity or cell-cell adhesion. To test this, we cultured MDCK II cells on semi-permeable supports for 14 days to generate polarized epithelial sheets, and performed ICC for NBCn1 and E-cadherin (Fig. 5A). Under control conditions, NBCn1 was exclusively localized to the basolateral membrane, with especially pronounced lateral expression (Fig. 5A, top panel). We next disrupted cadherin-based cell-cell adhesion by treating the epithelial layers with 3 mM EGTA for 2 h. Strikingly, in parallel with the expected internalization of E-cadherin (Nakagawa, 2001), this treatment caused NBCn1 to be almost completely internalized, remaining partially co-localized with internalized E-cadherin (Fig. 5A, middle panel). EGTA washout restored epithelial polarity within 4-6 h, and this correlated with near-complete re-localization of NBCn1 to the plasma membrane (Fig. 5A, lower panel). As a control, we carried out the same experiment for NHE1, which remained in the plasma membrane to a much greater extent than NBCn1 upon EGTA-mediated disruption of cell polarity (Suppl. Fig. 5).

**Figure 5.**
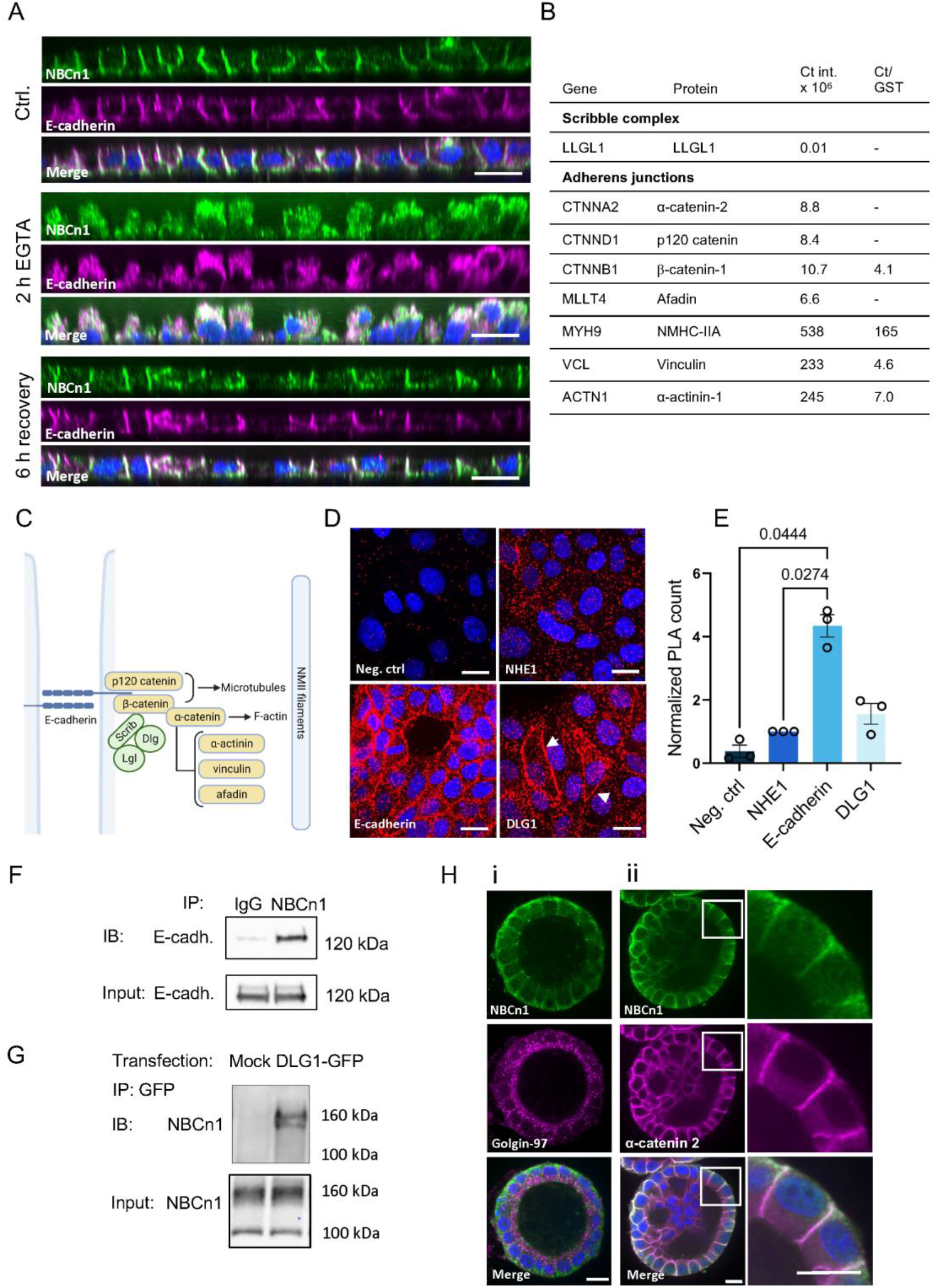
NBCn1 dynamically localizes to lateral domains and interacts with cell polarity and - adhesion proteins. MDCK II cells were grown for 14 days on transwells to polarize, then treated with 3 mM EGTA for 2 h followed by removal of EGTA and recovery in growth medium. Cells were fixed and stained with antibodies for NBCn1, E-cadherin and DAPI and imaged with z-stacks on a confocal microscope. **(A)** Representative side view images of non-treated cells (i), treated without recovery (ii), and cells recovered for 6 h (iii). **(B)** Putative binding candidates for the NBCn1 C-terminal in MCF-7 cells, based on MS analysis as described in (Olesen et al., 2018). The C-terminal/GST ratio was calculated based on intensities in the C-terminal pull-down and GST-pull-down. “–” indicates no intensity measured for GST. **(C)** Main components of the Scribble polarity complex and adherens junction proteins illustrating interactions between the MS hits shown in B. **(D)** Proximity ligation assay (PLA) of NBCn1 with E-cadherin and DLG1 in non-confluent MCF-7 cells. Negative control: only NBCn1 antibody. NHE1: control for plasma membrane proximity, not known to interact with NBCn1. Representative images with PLA signal in red, DAPI in blue. **(E)** Quantification of PLA signal (n=3), one-way ANOVA with Tukey’s multiple comparisons test. **(F)** Native co-IP in non-confluent MDCK II cells. Cells were lysed and subjected to pull-down with antibody against NBCn1 or isotype control (IgG) followed by western blotting of the pulldown and input fractions. Representative blots (n=3). **(G)** HEK293T cells were transiently transfected with DLG1-GFP and lysed. Lysates were subjected to pulldown against GFP followed by western blotting. Mock: transfected without plasmid. Representative blots (n=3). **(H)**: MDCK II cells were grown in Matrigel (i) or Geltrex (ii) for one week to form polarized cysts, fixed and stained with antibodies against NBCn1, Golgin-97, α-catenin 2, and DAPI. Scalebar 20 um.

To gain insight into the molecular mechanisms involved in this marked dependence of NBCn1 localization on epithelial polarity, we revisited our MS analysis (Olesen et al., 2018) and noted an abundance of lateral polarity- and cell-cell adhesion proteins among the putative NBCn1 interaction partners (Fig. 5B). These included polarity proteins of the Scribble complex (LLGL1), and multiple components of adherens junctions, including α-catenin-2, p120 catenin (CTNND1), β-catenin-1, afadin (aka MLLT4), non-muscle myosin-IIA (NMIIA, MYM9), vinculin and α-actinin (Fig. 5B) (Heuzé et al., 2019, Harris and Tepass, 2010). An overview of the organization of the main proteins in the two complexes is shown in Fig. 5C.

To validate the potential interaction of NBCn1 with these complexes in MDCK II cells, we performed *in situ* proximity ligation assay (PLA), which detects close proximity (< 40 nm) of two proteins in intact cells (Soderberg et al., 2006). To limit background reflecting the shared presence of the proteins in the plasma membrane, we included PLA of NBCn1 with NHE1 (with which it is not expected to physically interact beyond that they both localize to the plasma membrane) as an additional control, and normalized data to the NBCn1-NHE1 PLA signal (Fig. 5D-E). NBCn1 exhibited strong proximity labelling with E-cadherin - 4-fold higher than with NHE1 and ~11-fold higher than negative control (Fig. 5D-E), and efficiently pulled down native E-cadherin in co-IP experiments (Fig. 5F). The human Scribble complex comprises in addition to the Scribble protein itself also lethal (2) giant larvae 1 and 2 (LLGL1, −2), and discs large (DLG) (Bonello and Peifer, 2019). PLA signal for NBCn1 with DLG1 was also quantitatively higher than both controls and is seen very clearly in plasma membrane regions (Fig. 5D-E), and the interaction was confirmed by co-IP after DLG1 overexpression in HEK293 cells (Fig. 5G).

Finally, we asked whether the marked adherens junction-association of NBCn1 was also observed in 3D cysts, a preparation more closely mimicking native epithelia. MDCK II cells were cultured as 3D cysts, and co-localization of NBCn1 with α-catenin was investigated by confocal imaging (Fig. 5H). NBCn1 was basolaterally localized in the cysts, as shown in Fig. 5Hi, with the TGN marker Golgin-97 marking the apical region, and exhibited strong, mainly lateral, co-localization with α-catenin-2 (Fig 5Hii and zoom).

These results show that NBCn1 plasma membrane localization is strongly dependent on cell-cell adhesions, and that the transporter co-localizes and interacts physically with adherens junction and Scribble complex components.

### NBCn1 interacts with Rho GTPases and other F-actin regulators and localizes to leading edge protrusions

Upon remodelling of epithelial organization between front-rear- and apico-basal polarity, the Scribble complex is relocalized, and in front-rear polarized cells, the complex is found in the leading edge along with actin-regulatory proteins such as Rac1, Cdc42 (Rodriguez-Boulan and Macara, 2014, Nelson, 2009, Muthuswamy and Xue, 2012) and vinculin (Bays and DeMali, 2017). We have previously shown that NBCn1 localizes to the leading edge of migrating epithelial cells (Olesen et al., 2018), and the transporter was recently assigned a role in filopodia formation in endothelial cells (Boedtkjer et al., 2016). We therefore asked whether NBCn1 might, similar to its binding partners, move dynamically between adherens junctions and leading edge. In congruence with this notion, our MS data indicated strong interaction of NBCn1 with Rho GTPase family members Rac1, Cdc42, RhoA and RhoC, as well as with Arp2/3 and vasodilator-stimulated phosphoprotein (VASP), key regulators of lamellipodia- and filopodia formation (Fig. 6A). ICC analysis confirmed the strong colocalization of NBCn1, VASP, and F-actin in lamellipodia of non-confluent Hs578T breast cancer epithelial cells (Fig. 6B), and the partial but clear colocalization of NBCn1 with vinculin (Fig. 6C).

**Figure 6.**
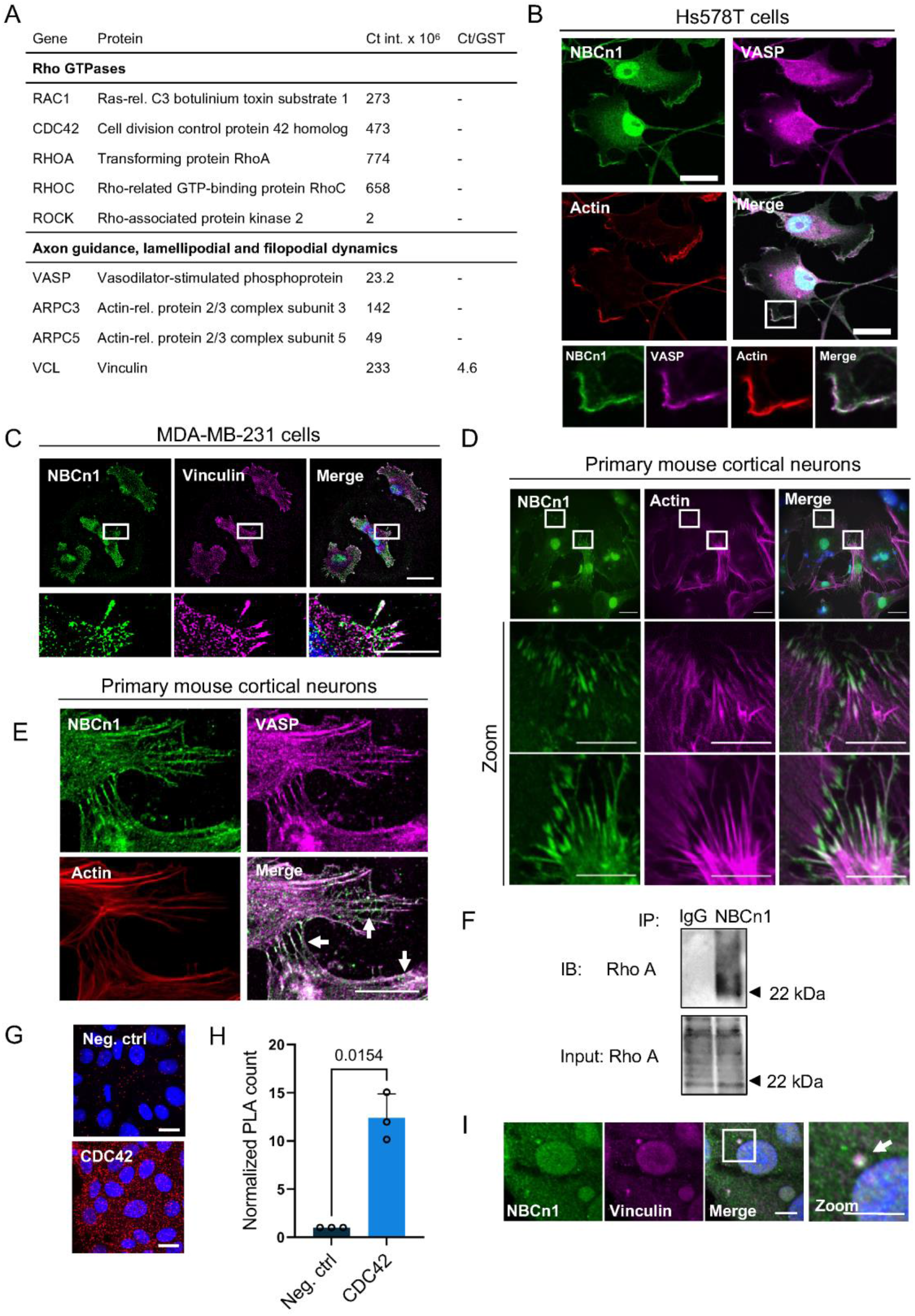
NBCn1 interacts with key regulators of cytoskeletal organization and dynamics. NBCn1 interaction partners involved in actin modulation, from MS analysis of NBCn1 C-terminal tail pulldown (Olesen et al., 2018). **(B)** Non-confluent Hs578T cells were fixed and stained for NBCn1, actin regulatory protein VASP and F-actin and imaged with fluorescence microscopy. Scale bar: 20 μm. (C) Non-confluent MDA-MB-231 cells were fixed and stained for NBCn1 and Vinculin. Scale bar: 20 μm. **(D-E)** Mouse primary cortical neurons were fixed and stained as in (B). Scale bar: 50 μm, zoom: 20 μm. **(F)** Native co-IP in non-confluent MDCK II cells. Cells were lysed and subjected to pull-down with antibody against NBCn1 or isotype control followed by western blotting of the pulldown and input fractions. Representative blots (n=3). **(G)** PLA with NBCn1 and Cdc42 in MCF-7 cells. Negative control: only NBCn1 antibody. Representative images with PLA signal in red, DAPI in blue. **(H)** Quantification of PLA signal (n=3). Statistical analysis: Paired, two-tailed Student’s t-test. **(I)** Non-confluent MCF-7 cells were fixed and stained for NBCn1 (C-terminal epitope antibody) and vinculin (n=3). Scale bar: 10 μm.

Vasp binds to the barbed end of actin filaments and is highly enriched at filopodial tips of neural growth cones (Lebrand et al., 2004). We therefore performed ICC of NBCn1 with F-actin and Vasp in primary murine cortical neurons (Fig. 6D-E) and observed a striking co-localization of the two proteins along actin filaments and filopodial tips. The interaction with Rho family GTPases was confirmed by co-IP of NBCn1 and RhoA (Fig. 6F) and by PLA of NBCn1 with Cdc42 (Fig. 6G-H).

Centrosomes were recently discovered to be actin-organizing centres, in which actin dynamics are regulated by Arp2/3 and other actin-regulatory proteins to coordinate centrosome positioning and spindle assembly (Farina et al., 2019, Inoue et al., 2019). Consistent with this, vinculin, a strong hit in our MS analysis (Fig. 5B, 6A) is a pericentriolar component (Chevrier et al., 1995), Indeed, ICC analysis revealed clear co-localization of NBCn1 and vinculin in perinuclear structures strongly reminiscent of centrosomes (Fig. 6I).

These data show that NBCn1 localizes to lamellipodia and filopodia and interacts with polarity-regulating proteins such as RhoA and Cdc42, and raise the possibility that NBCn1 might localize to centrosomes.

### NBCn1 localizes to centrosomes and spindle of epithelial cells

This unexpected finding led us to explore our MS data for hits relevant to such a localization. To our surprise, we found that multiple centrosomal proteins, including CEP135, CEP164, CEP290, and pericentriolar material protein 1 (PMC1), were strong putative NBCn1 interacting proteins (Suppl. Table 1). Furthermore, we noted strong pulldown of proteins involved in ciliary transport and ciliogenesis such as ADP-ribosylation factor-like protein-3 (ARL3) and intraflagellar transport protein 27 (IFT27); and proteins essential for mitotic spindle formation and –organization, such as microtubule-associated protein 4 (MAP4) and Ran-binding protein 1 (RANBP1) (Suppl. Table 1).

We therefore investigated whether NBCn1 might localize to these dynamic subcellular structures. Indeed, in non-confluent MDCK II cells co-stained for NBCn1 and the centrosomal marker γ-tubulin, we detected NBCn1 in the centrosome of a fraction of the non-dividing cells (Fig. 7A, Suppl. Fig. 6A-B). To avoid potential antibody- or fixation artefacts, we next co-expressed GFP-NBCn1 with the microtubule/centrosome marker End-Binding protein 3 (EB3)-tdTomato. Using live imaging, GFP-NBCn1 was clearly visible in the proximity of the nucleus of interphase cells, compatible with the location of centrosomes, and close to the poles and along the spindle of dividing cells (Fig. 7B, Suppl. Fig. 6F, Suppl. video 1). We also detected HA-tagged NBCn1 in the spindle of MDCK II cells using an HA-tag antibody (Suppl. Fig. 6C). Finally, we detected NBCn1 in centrosomes and spindle in multiple other cell types, using both N- and C-terminal epitope NBCn1 antibodies (Suppl. Fig. 6D). NBCn1 was also visible in the spindle of anaphase cells, as visualized by PLA in MCF-7 cells (Suppl. Fig 6E-F).

**Figure 7.**
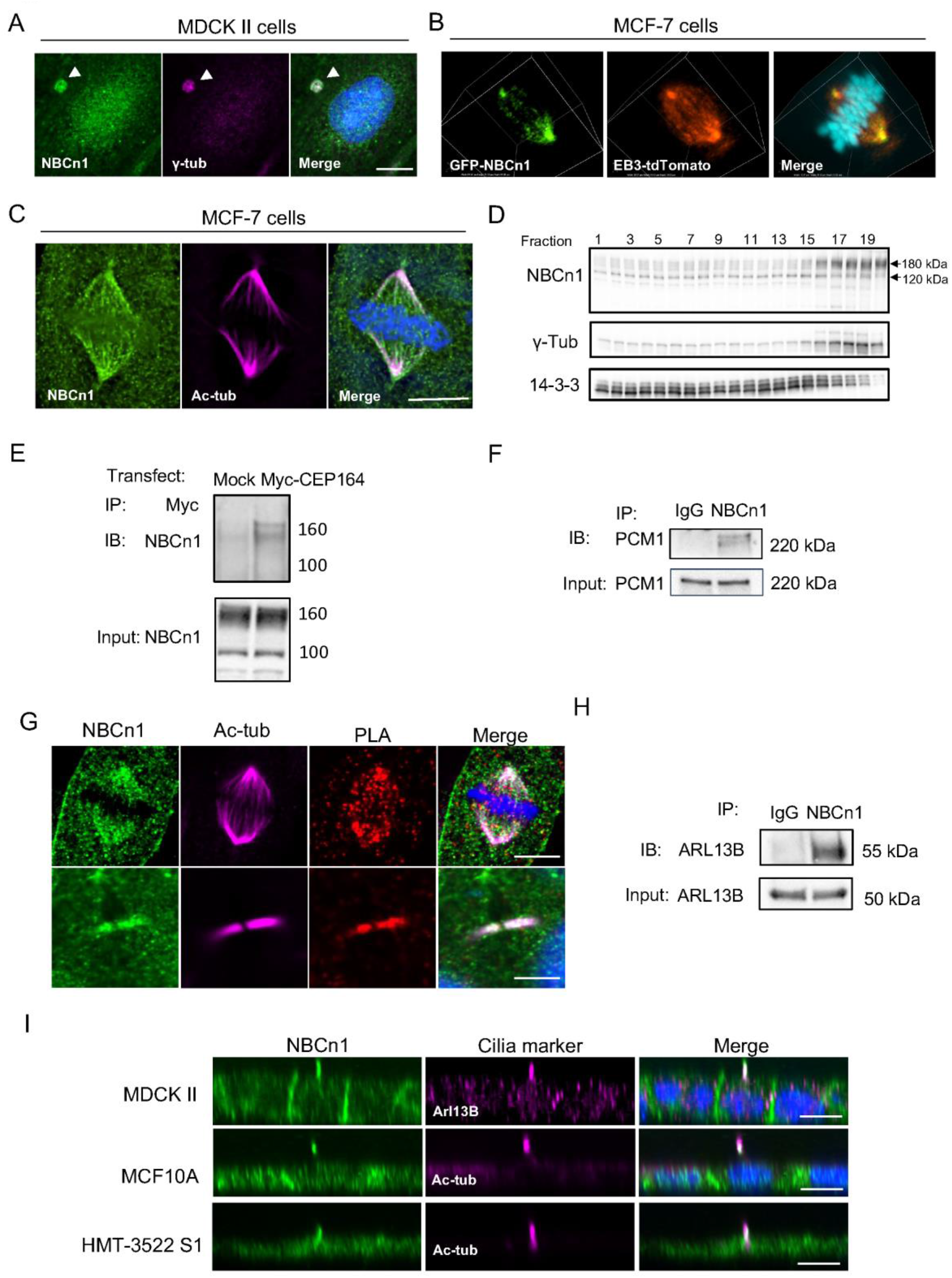
NBCn1 localizes to centrosomes, midbodies, cilia and the mitotic spindle. **(A)** Non-confluent MDCK II cells were fixed and stained for centrosomal marker γ-tubulin and NBCn1 (C-terminal epitope antibody).Scale bar: 10 μm. **(B)** MCF-7 cells were transiently double transfected with GFP-NBCn1 and centrosome and spindle marker EB3-tdTomato and Hoechst 33342, followed by live imaging by spinning disc confocal microscopy. Scale bar, 2 μM **(C)** MCF-7 cell from non-confluent culture, fixed and stained for NBCn1 (N-terminal epitope antibody) and acetylated tubulin (Ac-tub) to visualize spindles. A z-projected stack of three deconvoluted images is shown. Scale bar: 3 μm. **(D)** MCF-7 cells were treated with cytochalasin D and nocodazole to stabilize centrosomes. Cells were lysed and centrifuged on a sucrose cushion and fractions were blotted for NBCn1, centrosomal marker gamma tubulin and cytosolic marker 14-3-3. The same experiment was performed in RPE-1 cells with similar results (total n=4). **(E)** HEK293T cells were transiently transfected with Myc-CEP164, lysed and subjected to pulldown against Myc followed by western blotting for NBCn1. Mock: transfected without plasmid. Representative blots (n=3). **(F)** Native co-IP in non-confluent MCF-7 cells. Cells were lysed and subjected to pull-down with antibody against NBCn1 or isotype control (IgG) and western blotting of PCM1 in pulldown and input fractions. Representative blots (n=3) **(G)** Native co-IP as (F), blotted for Arl13B. **(H)** PLA for NBCn1 (C-terminal epitope antibody) and acetylated tubulin in non-confluent MCF-7 cells that were additionally fixed and stained for NBCn1 and acetylated tubulin to visualize spindles and midbodies (n=3). Scale bar: 10 μm. **(I)** MDCK II, MCF10A and HMT-3522 S1 cells were grown for 7 days on transwells to form polarized monolayers, fixed, and stained using NBCn1 C-terminal antibody (MDCK II and HMT-3522 S1) or NBCn1 N-terminal antibody (MCF10A) and ciliary markers Arl13B and acetylated tubulin. Imaged with z-stacks on a confocal microscope. Representative images seen from the side (n=3). Scale bar: 10 μm.

To further validate this unexpected finding, we isolated centrosomal fractions of MCF-7 cells and blotted them for NBCn1, centrosomal marker γ-tubulin and cytosolic marker 14-3-3. Non-centrosomal fractions showed roughly equal-intensity NBCn1 bands at ~120 kDa and ~180 kDa, consistent with the apparent MW of major NBCn1 splice variants (Parker and Boron, 2013). Strikingly, the 180 kDa band was strongly enriched in the centrosomal fraction (Fig. 7D). Co-IP of NBCn1 with the centrosomal proteins and MS hits, Centrosomal protein-164 (CEP164) and Pericentriolar material-1 (PCM1) (Suppl. Table 1), confirmed their interaction with NBCn1 (Fig. 7E-F). We reasoned that if NBCn1 was in the spindle and centrosome it might also be in midbodies - the structures that form from central spindle microtubules during cytokinesis and play key roles in regulation of postmitotic events (Peterman and Prekeris, 2019). Indeed, strong PLA signal for NBCn1 and tubulin was detected not only in the spindle but also in midbodies of MCF-7 cells (Fig. 7G).

These data provide evidence that NBCn1 localizes to the centrosome, spindle and midbodies of epithelial cells.

### NBCn1 localizes to primary cilia of polarized epithelial cells

After its resorption during mitosis, the primary cilium is re-assembled from the mother centriole of the centrosome during growth arrest (Christensen et al., 2008). Furthermore, the midbody has been reported to deliver material for the biogenesis of primary cilia in polarized epithelial cells (Labat-de-Hoz et al., 2020). We therefore asked whether NBCn1 could also be found in primary cilia. Consistent with this notion, NBCn1 pulled down the ciliary proteins Arl13B (Fig. 7H) and IFT27 (Suppl. Fig. 6G) in co-IP experiments, confirming their physical interaction. Next, we polarized three epithelial cell lines (MDCK II, MCF10A and HMT-3522 S1) on transwell filters for 7 days, followed by ICC analysis for NBCn1 and ciliary markers acetylated tubulin (acTub) or Arl13B. Clear NBCn1 localization was readily detectable along the length of the cilia in all three cell lines, using antibodies against N- or C-terminal epitopes of NBCn1 (Fig. 7I). Ciliary NBCn1 was detected in essentially all visible cilia in these cells, as seen in an overview of a polarized sheet of MDCK II cells stained for NBCn1 and Arl13B (Suppl. Fig. 6H). Importantly, in non-polarized Panc-1, mIMCD3, and RPE-1 cells that were serum depleted for 24 h to induce primary cilia formation, cilia were devoid of detectable NBCn1 (Suppl. Fig. 6I), suggesting that ciliary NBCn1 localization is restricted to polarized epithelial cells.

Taken together, these data show that NBCn1 localizes to primary cilia in polarized epithelial cells, yet is not detectable in primary cilia of non-polarized cells.

### NBCn1 is found in spindle-organizing Rab11-positive endosomes

While the presence of a membrane protein at spindle and midbody may at first glance seem surprising, several other examples of this are reported (Sauer et al., 2005, Skop et al., 2004), several studies have shown that the mitotic spindle is surrounded by membrane (Buch et al., 2009). But how might NBCn1 be trafficked to these subcellular locations, and what is the link between the localization of the transporter in spindle-midbody, centrosome, and primary cilium?

Notably, Rab11 was a strong potential interactor of the NBCn1 C-terminal in our MS analysis (Fig. 8A). Ciliogenesis occurs by two fundamentally different mechanisms, denoted the intracellular and alternative pathways (Labat-de-Hoz et al., 2020). The former involves delivery of components by Rab11-positive recycling endosomes. Rab11 endosomes are crucial for ciliary biogenesis (Knödler et al., 2010), interact with mother centriole appendages in interphase cells (Hehnly et al., 2012), traffic spindle assembly proteins to centrosomes and spindle for proper spindle alignment and cytokinesis (Hehnly and Doxsey, 2014, Hehnly et al., 2012). Illustrating their key role in cell division, Rab11 endosomes are involved in centrosome movement towards the cytokinetic bridge and necessary for cell abscission (Krishnan et al., 2022).

**Figure 8.**
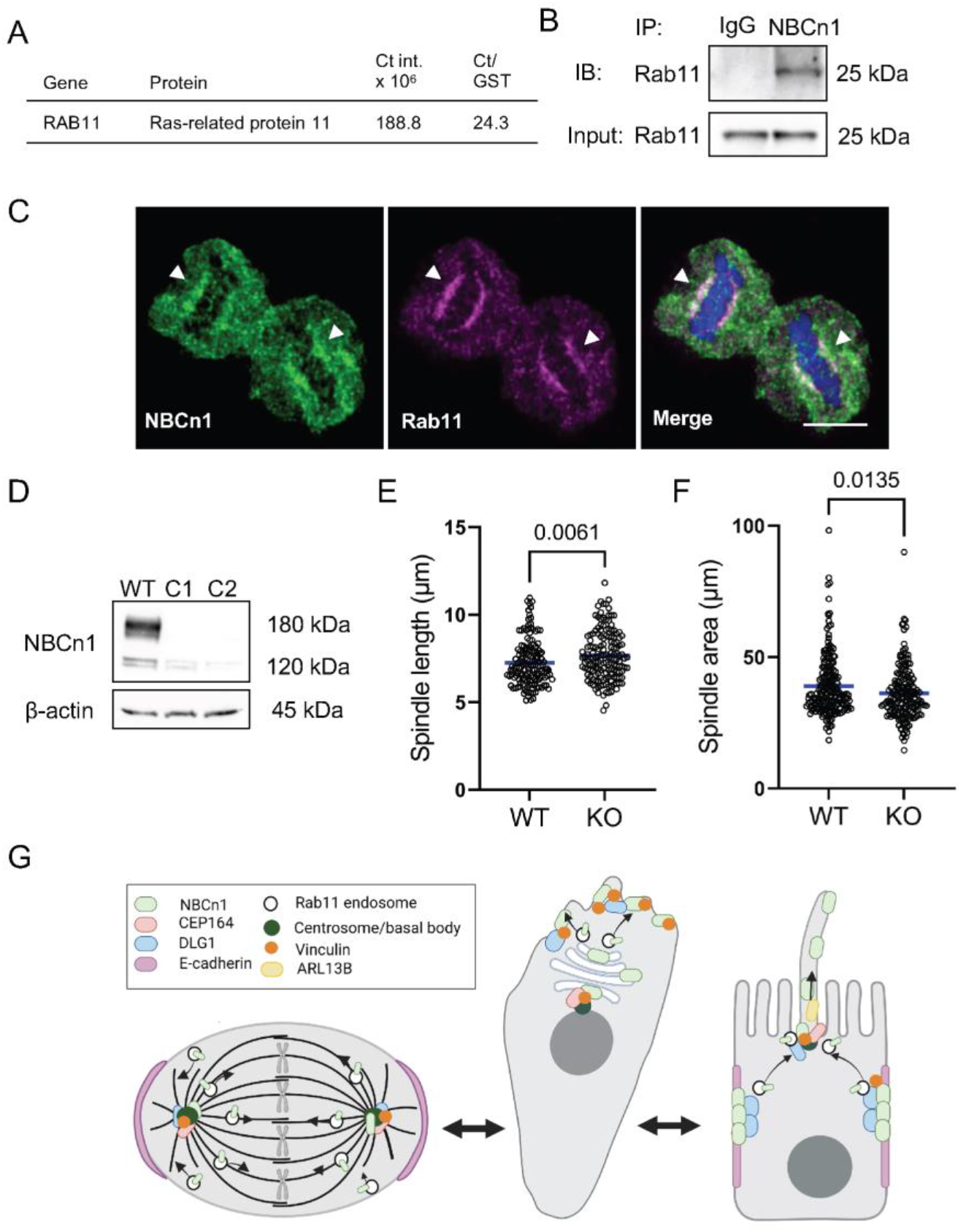
NBCn1 localizes to Rab11 endosomes and NBCn1 KO alters spindle area. **(A)** MS data shows Rab11 pulldown by NBCn1. From (Olesen et al., 2018). **(B)** Native co-IP was performed in non-confluent MDCK II cells. Cells were lysed and subjected to pull-down with antibody against NBCn1 or isotype control followed by western blotting of the pulldown and input fractions. Representative blots (n=3). **(C)** Non-confluent MCF-7 cells were fixed and stained for NBCn1 and Rab11. Scale bar: 10 μm. **(D)** Validation of NBCn1 KO in MDA-MB-231 cells. Spindle volume **(E)** and length **(F)** of MDA-MB-231 NBCn1 KO cells. Cells were grown for 24 h, fixed and stained for acetylated tubulin and centrosomal marker CEP164. Cells in metaphase were imaged with z-stacks and the largest spindle area in 2D measured as well as the length of the spindles pole-to-pole. Data represent all measurements from two clones, with 3-4 independent biological repeats each. Unpaired, two-tailed Student’s t-test. **(G)** Summary of findings and working model of NBCn1 trafficking with changes in epithelial cell polarity. See text for details.

We therefore pursued the possible role of Rab11 endosomes in regulating NBCn1 trafficking. Firstly, we showed that Rab11 was pulled down by NBCn1 in co-IP analysis (Fig. 8B), confirming our MS data. Next, we showed that NBCn1 and Rab11 co-localized to the mitotic spindle (Fig. 8C). Interestingly, transfection with dominant negative (DN) Rab11 caused the formation of Rab11 aggregates, in which NBCn1 clearly co-aggregated (Suppl. Fig. 7), suggesting that NBCn1 trafficking is compromised when Rab11 function is blocked.

The size of the mitotic spindle is determined by a series of microtubule-regulatory proteins as well as by cell size (Young et al., 2014). Microtubule assembly has been known for decades to be pH dependent (Regula et al., 1981), yet the impact of pH regulation on spindle size is to unelucidated. Strikingly, measurements of spindle area in WT and NBCn1 CRISPR/Cas9 knockout (KO) MDA-MB-231 cells (Fig. 8D) stained for acetylated tubulin revealed a significant increase in spindle length (Fig. 8D) and decrease in spindle area (Fig. 8E) in NBCn1 KO cells compared to WT cells.

Together, these findings show that NBCn1 co-localizes with Rab11 in the mitotic spindle, that interfering with Rab11 function causes partial NBCn1 mislocalization, and that depletion of NBCn1 appears to alter spindle morphology.

## DISCUSSION

The physiological function of ion transport proteins is entirely dependent on their subcellular localization. Local pH regulation by acid-base transporters is a case in point, having been shown to regulate cell differentiation, cell cycle progression, organelle function and -trafficking, motility, and signaling (Ulmschneider et al., 2016, Frantz et al., 2007, Pedersen et al., 2021, Chadwick et al., 2021). Furthermore, ion transport proteins can play local roles beyond transport, as scaffolds for signaling regulation (Hendus-Altenburger et al., 2016) or cell adhesion (Vagin et al., 2012); see (Meima et al., 2007). Here, we determine novel molecular determinants of NBCn1 plasma membrane trafficking, and we identify NBCn1 as a protein localized not only to the basolateral plasma membrane and cell-cell junctions, but also to primary cilia, centrosomes, mitotic spindle and midbodies.

### NBCn1 plasma membrane localization is dependent on N- and C-terminal folded motifs

We previously showed that NBCn1 plasma membrane localization is dependent on interaction of the proximal part of the NBCn1 C-tail with the scaffold protein RACK1 (Olesen et al., 2018). Here, we pinpoint a motif of only 6 amino acids (1133-1139) as essential for NBCn1 plasma membrane localization, with deletions of either the first or last 3 residues of the motif as equally deleterious. Furthermore, we demonstrate for the first time that both cytosolic domains of NBCn1 are required for its membrane transport: deletion of the first ~150 residues of the N-tail, in particular a predicted β-sheet region spanning residues ~100-150, also abolished plasma membrane localization. Given the very strong conservation of these regions across species and SLC4A family isoforms (Suppl. Fig. 1), we expect that the domains identified here are also important for membrane targeting of other isoforms. Specifically, the RELSWLD motif, which spans exactly the 6 essential amino acids in the NBCn1 C-tail is highly conserved, and identical in SLC4A7, −8, and −10 (Suppl. Fig. 1A-B).

Searching for physiological mechanisms regulating NBCn1 localization through these motifs we noted firstly that NBCn1 plasma membrane localization increased as cells underwent apico-basal polarization, was rapidly lost upon disruption of cadherin-dependent cell-cell interactions, and rapidly reversed upon their recovery. Second, NBCn1 interacted strongly with E-cadherin and DLG1, interestingly consistent with adherens junction- and polarity protein interactions indicated by recent BioID data (Go et al., 2021). We therefore propose that plasma membrane localization of NBCn1 in epithelial cells is dependent on targeting and retention of the transporter to adherens junctions and the scribble apical polarity complex as these become established. Cell-cell adhesion dependence is also demonstrated for the Na^+^,K^+^ ATPase (Vagin et al., 2012), yet does not appear to be generic for basolateral transporters as another pH regulator, NHE1, was much less cell-cell contact dependent. Notably, NBCn1 exists in a striking number of splice variants, which differ between cell types (Parker and Boron, 2013), and future works should explore the targeting mechanisms in non-epithelial cells.

We found that steady state pH_i_ was lower in fully polarized than in sparsely seeded cells. Interestingly, this is consistent with elegant work by Murer and colleagues (Vilella et al., 1992) showing that because of the tightness of the cell-cell junctions and basolateral localization of acid-base transporters, polarized MDCK cells are unable to recover pH_i_ after acidification unless they are basolaterally perfused. However, the lower pH_i_ is not the cause of the strong NBCn1 plasma membrane localization in polarized cells, as NBCn1 membrane localization was unaltered if pH_i_ was instead reduced in non-polarized cells by reducing pH_e_ (Michl et al., 2022). In contrast, NBCn1 plasma membrane localization was increased by elevation of cytosolic cAMP levels, similar to what was reported for NBCe1 (Yu et al., 2009, Carranza et al., 1998). This does not exclude additional driving mechanisms but it seems possible that the establishment of the lateral complexes may in itself be sufficient for the shift in NBCn1 membrane localization upon polarization.

### NBCn1 localizes strongly to filopodia and other leading edge actin structures, interacting with vinculin and Rho family GTPases

Transition between apico-basal and front-rear polarity involves extensive reorientation of the Golgi apparatus, centrosome, and the associated microtubule network. During migration, Scribble, DLG-1 and LGL localize to the leading edge where they regulate front-rear polarity and migration in interplay with Cdc42, Rac, and other polarity proteins and cytoskeletal regulators (Muthuswamy and Xue, 2012, Rodriguez-Boulan and Macara, 2014, Nelson, 2009). We show here that NBCn1 interacts with Cdc42 and is strongly enriched in leading edge filopodia and lamellipodia, co-localizing with vinculin and VASP, interactions supported both by our mass spec data and by BioID studies (Go et al., 2021, Bagci et al., 2020). The possible role of NBCn1 in cell migration is unclear, with a role in initial migration rates reported in endothelial cells (Boedtkjer et al., 2016), whereas in breast cancer cells (i.e. epithelial-derived cells), such an effect was not detectable despite strong NBCn1 expression (Lauritzen et al., 2012). We therefore suggest that the pronounced enrichment of NBCn1 in leading edge structures could contribute to polarization rather than migration *per se*, through local regulation of pH sensitive elements of the polarization process such as Cdc42 (Frantz et al., 2007). Interestingly, Cdc42 regulates microtubule-plus end organization through adenomatous polyposis coli (APC) regulation at microtubule plus-ends and formation of DLG1 puncta in the leading edge membrane (Etienne-Manneville et al., 2005). The tight interaction of NBCn1 and DLG1 would put NBCn1 in a prime position to regulate the pH sensitive process of microtubule elongation (Regula et al., 1981) and NBCn1 interaction with VASP and Arp2/3 could in a similar manner contribute to formation of leading edge filopodia and lamellipodia.

### NBCn1 is present in primary cilia, centrosomes, spindle, and midbodies

A key discovery in this study is that NBCn1 is found in primary cilia, centrosomes, mitotic spindle, and midbodies. Of these, the ciliary localization is the least surprising. Cilia are membrane-bound organelles and are well known to express functional ion channels such as the widely studied polycystin-2, the transient receptor potential channel TRPC1, and the epithelial sodium channel αENaC (Raychowdhury et al., 2005). Also in agreement with our findings, a ciliary membrane proteome included NBCn1 (Kohli et al., 2017). However, our work is the first report of a pH regulatory protein in the primary cilium and raises the question of how ciliary pH is regulated, something that has to our knowledge never been explored.

Interestingly, despite the prominent ciliary NBCn1 localization in polarized epithelia, we could not detect the transporter in primary cilia of non-polarized cells. This suggests that mature epithelial cell-cell adhesions are important for trafficking of NBCn1 to the primary cilium. The mechanisms targeting membrane proteins to primary cilia are remarkably diverse, and both direct trafficking and docking to plasma membrane regions, followed by lateral movement into the cilium, have been proposed (Morthorst et al., 2022, Monis et al., 2017, Nachury, 2018). Notably, the strong NBCn1 interaction partner DLG1 is a well-established binding partner of the kinesin-3 family motor protein KIF13B. Interaction of DLG1 with KIF13B initiates KIF13B-mediated DLG1 transport along microtubules and is involved in the trafficking of the membrane protein SMO to the ciliary membrane (Morthorst et al., 2022). We therefore speculate that NBCn1 delivery to the primary cilium could be dependent on its interaction with DLG1 followed by KIF13B-mediated delivery to the ciliary base in Rab11 endosomes (see below and Fig. 8E).

The discovery that NBCn1 also localized to the centrosome, mitotic spindle and midbodies of epithelial cells was fully unexpected, but localization of transport proteins to these structures is in fact not without precedent. The monocarboxylate transporter MCT4 has been found in the HeLa cell spindle (Sauer et al., 2005), localization of the glucose transporters GLUT1 and −4 to midbodies was required for abscission in CHO cells (Skop et al., 2004). The local function of these transporters is to our knowledge unexplored. However, in a remarkable parallel to our findings, the voltage gated potassium channel Kv10.1 was shown to localize to the centrosome where it modulates spindle dynamics via ORAI1 (Movsisyan and Pardo, 2020) and to the primary cilium, where it induces ciliary disassembly at onset of mitosis (Sanchez et al., 2016). As discussed above for the leading edge, NBCn1 could tune spindle organization through local pH regulation. This would be consistent with the major importance of NBCn1 for cell cycle progression (Flinck et al., 2018), and our finding of altered spindle area in NBCn1 KO cells points in this direction.

### A possible model of dynamic NBCn1 trafficking between subcellular compartments

How can we understand the localization of a transmembrane protein such as NBCn1 to all of these compartments? Firstly, all of these structures contain membranes, as recently demonstrated for the localization of the integral nuclear membrane protein Samp1 to the polar regions of the mitotic spindle (Larsson et al., 2018). Recent evidence has made it clear that lateral adhesion and polarity complexes, spindle/midbodies, centrosomes and leading edge are closely and dynamically linked. DLG1 is an important regulator of spindle polarity and movement (Porter et al., 2019), as well as of front-rear polarity in migrating cells, as noted above (Muthuswamy and Xue, 2012, Rodriguez-Boulan and Macara, 2014, Nelson, 2009). Centrosomes are now recognized to be actin organizing centres, to which numerous focal adhesion and actin regulatory proteins localize, including NBCn1 interactors vinculin (Chevrier et al., 1995), ARP2/3 (Farina et al., 2019), and DLG1 (van Ree et al., 2016). Thus, we propose that NBCn1 is part of a dynamic protein interactome which translocates between basolateral adherens junction/Scribble complex region, leading edge structures, centrosomes, spindle, and the spindle remnant, the midbody (Fig. 8G). When cells are apico-basally polarized, NBCn1 may translocate from the basolateral region to the ciliary base in Rab11-containing endosomes (see below). Also consistent with this, Rab8a, which is involved in ciliogenesis through NBCn1 interactors CEP164 and CEP290 (Tsang et al., 2008, Schmidt et al., 2012), was another hit in our MS analysis. Not mutually exclusive with this, NBCn1 may also traffic to the cilium via its localization in the midbodies, in the alternative pathway of ciliary biogenesis, in which midbodies provide material for the growing cilium (Labat-de-Hoz et al., 2020). This has been found to be the pathway employed in MDCK II cells (Bernabé-Rubio et al., 2016), and would, interestingly, be consistent with our finding that ciliary NBCn1 is only detected in cilia of polarized epithelial cells.

The precise molecular mechanisms enabling such dynamic relocation of NBCn1 with the polarization states of the epithelial cell - mitotic, front-rear, and apico-basal – remain to be uncovered. Again, Rab11 seems to be a possible link. Recent work points to a key role for Rab11 endosomes in trafficking to interphase centrosomes and mitotic spindle (Zhang et al., 2008). Rab11 endosomes were found to decorate centrosomes as well as mitotic spindles and contained crucial spindle assembly factors in much higher concentrations during mitosis than in interphase (Hehnly and Doxsey, 2014). Intriguingly, a similar function of the *C. elegans* Rab11 homolog RAB-11 was dependent on RACK1 for its localization to the mitotic spindle and centrosomes (Ai et al., 2009), and RACK1 also localizes to midbodies in mammalian cells (Skop et al., 2004). It is tempting to suggest that the interaction between RACK1 and NBCn1 which we previously demonstrated (Olesen et al., 2018) could contribute to directing NBCn1 to Rab11 endosomes, via which it would be trafficked to centrosomes, spindle, midbodies, and primary cilia and contribute to local pH regulation. While the precise functional role of NBCn1 is in these localizations remains to be elucidated, we note that the spindle area/length parameters are altered upon NBCn1 KO, suggesting a role for local NBCn1 transporter activity in spindle assembly and morphology. This has implications for similar transporters that may be playing hitherto unexplored intracellular roles, and this will be an exciting question for future research.

*In conclusion*, we show here that both N- and C-terminal motifs in NBCn1 are essential for its plasma membrane targeting, and we map the C-terminal motif to a six proximal amino acids. Further, we provide detailed evidence that NBCn1 also localizes to centrosomes, spindle, midbody and primary cilia, cycling between these compartments as cell polarity changes.

## MATERIALS AND METHODS

### Antibodies and reagents

A list of all antibodies used is provided in the online Supplementary Materials. Other reagents appear below in the order of use.

### Cell culture and treatments

MCF-7 cells were cultured in DMEM 1885 supplemented with 5% fetal bovine serum (FBS), 1% penicillin/streptomycin, and 1% non-essential amino acids. NBCn1 KD and plKO.1 control cells were additionally cultured with 1 μg/mL Puromycin. MDCK II cells were cultured in alpha modified MEM (α-MEM) with 10% FBS, 1% penicillin/streptomycin (P/S) and 2 mM L-Glutamine. MDA-MB-231 cells were cultured in DMEM 1885 with 10% (FBS), 1% P/S and 1% non-essential amino acids. HEK293T cells were cultured in DMEM (Gibco, #41966-029) with 5 % FBS, 1% P/S, and 1% non-essential amino acids. mIMCD3 cells were cultured in DMEM (Gibco, #11995-065) 1:1 with F12 Ham (Sigma, #N6658),10% FBS and 1% P/S. RPE-1 and Hs578T cells were cultured in DMEM (Gibco, #41966-029) with 10% FBS and 1% P/S. MCF10A cells were cultured in DMEM (Gibco, #41966) 1:1 with Ham’s F12 nutrient mixture medium (Sigma, #N6658), 5% FBS, 1% P/S, 20 ng/ml recombinant human epidermal growth factor (hEGF, #E9644, Sigma), 0.25 ng/mL hydrocortisone (#H0888, Sigma), and 10 μg/ml bovine insulin-transferrin-selenium (Gibco, #41400-045). Panc-1 cells were grown in DMEM (Gibco, #32430-027) with 10% FBS and 1% P/S. HMT-3522 S1 cells were grown in DMEM (Gibco, #41966) 1:1 with Ham’s F12 nutrient mixture medium (Sigma, #N6658) with 2 mM glutamine, 250 ng/ml insulin (Sigma, #I6634), 10 μg/ml transferrin (Sigma, #T2252), 10 nM sodium selenite (Sigma, #S9133), 0.1 nM 17 beta-estradiol (Sigma, #E2758), 0.5 μg/ml hydrocortisone (Sigma, #H0888), 5 μg/mL bovine prolactin and 10 ng/ml hEGF. Recombinantly expressed human prolactin was a kind gift from Prof. B. Kragelund, Dept. of Biology, University of Copenhagen.

All cell lines were maintained at 37°C, 95% humidity and 5% CO_2_. Where indicated, cells were grown for 7-14 days on Transwell polycarbonate filters (Corning Incorporated (Costar), Sigma-Aldrich, #3401) with 0.4 μm pore size.

### Constructs and transfections

GFP-constructs: Rat NBCn1 (rNBCn1-D, NM_058211.2) was amplified by PCR from cDNA in plasmid kindly provided by Dr. Ebbe Boedtkjer (Aarhus University, Denmark) and inserted into pEGFP C1 and N1 vectors to produce C1 as previously described (Olesen et al., 2018) to produce the N and C-terminally GFP-tagged rNBCn1-D, respectively. Eight constructs with GFP-NBCn1 were produced including full length and ΔCt-NBCn1 described in (Olesen et al., 2018). In addition, Δ1139-1254, Δ1133-1254, Δ1133-1139, Δ1133-1135, Δ1136-1139, F1130A were produced. Five NBCn1-GFP constructs were produced: Full length, K135A, K142A, Δ1-98 and Δ1-148. Deletion and point mutation constructs were produced through inverse PCR using Takara In-Fusion HD cloning kit (#638920) (Takara Bio Inc., Kusatsu, Shiga, Japan). Primers used are listed in supplementary materials.

Plasmids were transiently transfected into HEK293T or MCF-7 cells at 70% confluence using Lipofectamine 3000 (Invitrogen, #L3000015) in a 4:1 v/w ratio following manufacturer’s protocol. Medium was replaced with fresh growth medium after 8 h incubation with transfection mixture. Twenty-four h after transfection cells were washed in PBS and subjected to either immunofluorescence or co-IP assays. The following plasmids were employed: pEGFP-N1 (GenBank: U55762.1), pEGFP-C1(GenBank: U55763.1) DsRed-Rab11 dominant negative (DN) (Addgene: 12680), DsRed-Rab11 WT (Addgene: 12679), DLG1-GFP (Wu et al., 1998), CEP164-Myc (Addgene: 41148), and EB3 tdTomato ((Merriam et al., 2013), Addgene #50708).

### Fluorescence-based NBCn1 localization assay

MCF-7 cells were cultured on 12-mm glass coverslips, transiently transfected with NBCn1 constructs as described above, and fixed post-transfection with 2% paraformaldehyde for 20 min at room temperature. Nuclei were stained with DAPI, and cells were mounted on microscope slides and imaged on an Olympus IX83 microscope using cellSens dimension software. Quantification of NBCn1 localization was done by blinded analysis of a minimum of 50 cells per construct per experiment and classifying localization as plasma membrane, organellar, or a mixture of the two. Three researchers independently obtained almost identical results when counting the same experiments blinded as an extra layer of control.

### Immunocytochemistry (ICC) – general protocol

Cells were washed in cold PBS and fixed with 2% paraformaldehyde (Sigma, #47608) diluted in phosphate buffered saline for 15 min at room temperature, followed by 30 min on ice then washed 3×5 min in PBS. Cells were then permeabilized with 0.5% Triton-X-100 in TBST for 5 min and blocked with 5% BSA in TBST for 30 min. Coverslips were incubated with primary antibodies diluted in 1% BSA in TBST overnight at 4°C and with fluorophore conjugated secondary antibodies for 1-2 h at room temperature.

### Analysis of confluence-dependent localization

MDCK II cells were cultured on 12 mm glass coverslips for 8 days to a fully confluent, polarized cell layer, or for 24 h to reach 20/60% confluence. Cells were washed in PBS and fixed with 2% PFA for 15 min and subjected to the ICC protocol described above. Coverslips were mounted on microscope slides and images acquired with an Olympus IX83 microscope using cellSens Dimension Software. Data were quantified using line scan analysis as described below.

### Analysis of stimulus-dependence of localization

MDCK II cells were cultured on 12 mm glass coverslips and seeded the day before treatments with cells having reached 15-40% confluence at the time of stimulation. Growth medium with 20 μM HCl, 20 μM forskolin or 1 μg/mL EGF was added to cells after sterile filtration via a 0.22 μm filter (Millex-GP, #SLGP033RB). Control cells were provided with fresh growth medium and when indicated vehicle (DMSO, Sigma, #D2650) was added. Forskolin (#F6886, stock concentration: 10 mM diluted in DMSO) and epidermal growth factor (EGF, #E9644, stock concentration: 40 μg/mL in 10 mM acetic acid + 0.1% BSA) was from Sigma Aldrich, St. Louis, USA. Hydrochloric acid (HCl) was kept in a stock concentration of 1 M in ddH2O. Cells were fixed 24 h post-stimulation and subjected to IF analysis as above.

### Quantification via line scan analysis

Quantification of relative plasma membrane localization was performed by line scan analysis using ImageJ software. A line selection was drawn 90° across the membrane using E-cadherin staining as a membrane marker. The intensity of NBCn1/NHE1 staining along the line selection was measured as a ratio between the membrane intensity (peak intensity of E-cadherin + 3 adjacent values) and the cytosolic intensity (remaining values with a minimum of 30 values). For each line selection two intensity ratios were calculated, one for each of the two neighbouring cells.

### SDS-PAGE and Western blotting

Protein concentrations were determined by DC assay (BioRad, Hercules, USA). Samples with equal amounts of protein were mixed with a 1:1 mixture of 4x NuPAGE LDS Sample Buffer (Invitrogen, #NP0007) and 500 mM DTT (Sigma, #646563). Proteins were denatured for 5 min at 95°C and separated by SDS-PAGE using precast 10% polyacrylamide 10-well (Bio-Rad, #456-1034) or 18-well (Bio-Rad, #567-1034) gels, premade Tris/Glycine/SDS electrophoresis buffer (BioRad, #161-0732), and BenchMark protein ladder (Invitrogen, #10747-012). Proteins were transferred by Trans-Blot Turbo transfer system (BioRad) to a nitrocellulose membrane (BioRad, #170-4158/#170-4159), stained with Ponceau S (Sigma, #7170), and blocked in 5% non-fat dry milk in TRIS-buffered saline with Tween (TBST; 0.01 M Tris/HCL, 0.12 M NaCl, 0.1% Tween 20, pH 7.5) for min. 1 h at 37°C. Membranes were incubated overnight at 4°C with primary antibodies diluted either in 5% non-fat dry milk in TBST or 5% Bovine Serum Albumin (BSA, Sigma, # A7906) + 0.02% sodium azide in TBST. Next day, membranes were incubated with horseradish peroxidase (HRP) conjugated secondary antibodies diluted in 5% non-fat dry milk in TBST for 1 h at room temperature. Bands were visualized by chemiluminescence using enhanced chemiluminescent (ECL) substrate (BioRad, #170-5061/Cell Signaling, #12757S).

### pH measurements

MDCK II cells were cultured in WillCo-dishes (HBST-3522) and seeded either 7 days before pH measurements to reach 100% - or 24 h before to reach 20% - confluency. 4 μM of SNARF™-5F 5-(and-6)-carboxyl acid, AM ester, acetate (Invitrogen, S23923) was added to the cells and incubated for 30 min at 37 °C, 95% humidity and 5% CO2. The cells were then washed 3 times in MDCK II culture medium to remove excess SNARF and incubated again at 37 °C, 95% humidity and 5% CO_2_ for the cells to activate the SNARF. Immediately before the pH measurements the medium was changed to a Ringer solution (in mM: NaCl 115, KCl 5, Na_2_HPO_4_ 1, CaCl_2_ 1, MgCl_2_ 0.5, NaHCO_3_ 24), and placed on the microscope stage, heated to 37°C. For the pH measurements an addition of humid air with 5% CO_2_, 19.5% O_2_ and 75.5% N_2_ was always present. SNARF fluorescence was measured at 580 and 660 nm emission after excitation at 488 nm using an epifluorescence microscope (Nikon Eclipse Ti2). Steady state pH was measured for 5 min. The ratio of the intensity at 580 and 660 nm was calculated and converted to pH values using *in situ* calibration. The calibration curve was created using a KCl Ringer with 10 μM Nigericin (in mM: KCl 140, K_2_HPO_4_ 1, CaCl_2_ 1, MgCl_2_ 0.5 and a 1:1:1 mix of MOPS-TES-HEPES - 25) at the pH values of 6.2, 6.6, 7.0, 7.4 and 7.8.

### Transwell assays

For 2D polarized epithelial experiments, 25.000 cells were seeded in 0.4 μm transwell chambers (Corning Costar, #3401) and grown for 1-2 weeks as indicated. Medium was changed every 3 days and membranes were fixed and excised and treated as described for ICC general protocol.

### Disruption of cell-cell adhesion

Cells were washed once in Ca^2+^ free 3 mM EGTA Ringer and incubated for 2 h in fresh Ca^2+^ free 3 mM EGTA Ringer, followed by two washes in growth medium and incubation in growth medium for 0.5, 1, 2, 4, and 6 h. At these time-points, cells were washed in cold PBS and fixed as described for ICC general protocol.

### Preparation of MDCK II cysts

To form cysts, cells were grown in Nunc Lab-Tek II 8 well Chamber Slide System (Thermo Scientific, #154534) using either Matrigel (Corning, #356231) or Geltrex (Thermo Scientific, #A1413201). For experiments with Geltrex: The chambers were prepared by distributing 60 μl Geltrex to the bottom of each well followed by 20 min incubation at 37°C to polymerize the gel. MDCK II cell suspensions were prepared in cold medium containing 25 μl/ml Geltrex and seeded at 10,000 cells per 400 μl in the chamber slide wells. For experiments with Matrigel: 100 μl cold cell suspension containing 10.000 cells was gently mixed with 100 μl Matrigel (10 mg/ml) in Eppendorf tubes and 200 μl was immediately seeded into each well, incubated 20 min at 37°C to polymerise, and 300 μl growth medium was added on top. For both experiments, cells were cultured for one week, with growth medium replaced every 2 days. All steps for culturing and IF were performed by carefully pipetting with a 1000 μl pipette at the corner of the wells. For IF analysis of cysts, cells were treated in the chambers. This protocol was adapted from (Giles et al., 2014). Cells were washed thrice with DPBS containing Ca^2+^ and Mg^2+^, 200 μl 4% PFA in PBS was added to each well and slides were placed on a shaker at low speed for 30 min at room temperature (RT) to dissolve the gel and allow the cysts to stick to the glass slides. Cysts were washed 5 × 5 min in DPBS, incubated for 30 min at RT in permeabilization buffer (5% BSA in DPBS with 0.5% Triton-X), and with primary antibodies in permeabilization buffer over night at 4°C. Cells were washed 3×10 min in permeabilization buffer, incubated for 1-2 h at RT with fluorophore conjugated secondary antibodies in permeabilization buffer, washed, incubation with DAPI for 10 min, and washed 3 × 5 min in PBS. The top chamber walls were carefully removed, and the spheroids were mounted with N-propyl-gallate. Two coverslips were added on top of the cells and sealed with nail polish. Cysts were visualized using an inverted Olympus Cell Vivo IX83 with a Yokogawa CSU-W1 confocal scanning unit.

### Proximity ligation assay

Cells were fixed and permeabilized as described for IF analysis. The assays were carried out using Duolink PLA kit (Sigma-Aldrich, #DUO92002) following manufacturer’s protocol. Briefly, blocking solution was added for 60 min at 37° C. Coverslips were then incubated with primary antibodies overnight at 5° C. Each incubation step was followed by washing 3×5 min in TBST. PLA probes (PLUS and MINUS) were added and incubated for 1 h at 37°C. Ligase solution was then added and incubated for 30 min at 37°C followed by incubation with amplification solution containing polymerase for 100 min at 37°C and protected from light. In some experiments, secondary fluorophore conjugated antibodies were added to image co-localization of proteins with PLA dots. Coverslips were incubated with DAPI for 5 min at RT, followed by washing in PBS. Coverslips were then mounted with N-propyl-gallate, sealed, and visualized using an inverted Olympus Cell Vivo IX83 with a Yokogawa CSU-W1 confocal scanning unit. For quantification experiments, coverslip identity was blinded for the observer. Images were quantified with ImageJ using the Analyze Particles function on binary images with background subtraction. Minimum 150 cells were quantified per condition per experiment.

### Native co-immunoprecipitation

Cells were grown for either 7-10 days (confluent cells) or 24 h (non-confluent cells) in 10 cm Petri dishes, washed in ice-cold PBS and lysed in room temperature NP40 lysis buffer (50 mM Tris pH 7.4, 140 mM NaCl, 3 Na_3_VO_4_, 1% v/v IGEPAL CA-360 (Sigma, #I8896), Phosphatase Inhibitor cocktail (PhosStop) and Protease Inhibitor mix (cOmplete™) (Roche, #04906845001 and #1169748001). Lysates were homogenized using a 0.5 mm syringe needle and normalized to a protein concentration of 4 mg/ml in 500 μl. Samples were incubated overnight at 4°C with primary antibodies rotating end-over-end. For equilibration, Dynabeads Protein G (Invitrogen, #10004D) were washed 2x 10 min at 4°C with lysis buffer while gently rotating and immune complexes were incubated with 1.5 mg washed Dynabeads for 45 min at 4°C rotating end-over-end. Dynabeads with bound protein were washed 5×5 min in lysis buffer, boiled 5 min at 95 °C in 80 μl NuPAGE LDS Sample Buffer (Invitrogen, #NP0007) and dithiothreitol, vortexed thoroughly, and placed on ice for 30 min. Eluted protein complexes were separated with SDS-PAGE and analysed by Western blotting as described above.

### Cold buffer treatment for Rab11-NBCn1 co-localization experiments

Coverslips were washed twice in PBS containing calcium and magnesium, then placed in a cold Ringer buffer on ice for 20 min to preserve microtubule kinetochore fibres. The cold buffer contained 138 mM NaCl, 5 mM KCl, 1 mM Na_2_HPO_4_, 1 mM CaCl_2_, 0.5 mM MgCl_2_ and 20 mM Hepes at pH 7.4. Cells were then washed and subjected to the immunofluorescence protocol described above.

### Centrosome purification

Centrosome purification was performed as in (Sanchez et al., 2016). Briefly, cells were treated with cytochalasin D and nocodazole to stabilize centrosomes. Cells were lysed, intact cells and gross debris were removed through a 60% sucrose cushion and the cleared lysate was centrifuged on a discontinuous sucrose gradient (40-70%) at 120,000 *xg* for 60 min at 4 °C in a swinging bucket rotor. 200 μL fractions were collected and blotted for NBCn1, centrosomal marker gamma tubulin and cytosolic marker 14-3-3.

### Live imaging

Cells were cultured and transfected in four-well microscopy dishes (Ibidi #80426 μ-Slide #1.5 polymer coverslip). Images of living cells were acquired using a Nikon TiE-Andor spinning disk microscope (Nikon) with environmental control chamber and an Andor iXon Ultra 897 EMCCD camera, and using a 100 X oild immersion objective (Nikon PLAN APO-TIRF 100X 1.49 N.A.). Transfected cells were incubated with 100 nM Hoechst 33342 for 10 minutes, washed with imaging solution (in mM: 140 NaCl, 2.5 KCl, 1.8 CaCl_2_, 1 MgCl_2_, 20 Hepes, 20 glucose, pH 7.4) and immediately transferred to the microscope. Acquisition was controlled using NiS-Elements software. Planes for stacks were acquired at 0.2 μm intervals, and Imaris software (Oxford Instruments, v. 9.7) was used for 3D reconstruction.

### CRISPR/Cas9 knockout of NBCn1 in MDA-MB-231 cells

Briefly, the guide RNA (gRNA) sequence gRNA sequence CCAGCATGACTGTTCCATTG (exon 5) of NBCn1 was designed using CRISPR Finder software (https://wge.stemcell.sanger.ac.uk/find_crisprs), cloned into the pSpCas9(BB)-2A-Puro plasmid (Addgene #62988) and transfected into MDA-MB-231 cells using Lipofectamine^®^ 3000 (Invitrogen) according to the manufacturer’s protocol, retransfecting after ~3 days to increase delivery. Editing efficiency was assessed using the EnGen Mutation Detection Kit (NEB) (E3321S) according to the manufacturer’s protocol. Following single cell clone isolation and expansion, NBCn1 KO was verified by western blotting.

### Spindle measurements

Spindle length and area was measured in metaphase cells. Spindles were imaged in their central plane in 2D. Length was measured pole-to-pole while area was measured by overlaying an ellipse fitted to the spindle shape. The observer was blinded.

### Statistics

Data is shown as representative images from individual experiments or as quantitative analysis of several experiments. Plasma membrane localization, WB and PLA: one-way analysis of variance (ANOVA) with Tukey’s post-test. Line scan analyses: Fig. 3: one-way ANOVA (3 groups) with Dunnett’s multiple comparisons test. Fig 4 and supplementary Fig 3: non-parametric Mann Whitney U test (2 groups). Spindle and pH measurements: unpaired t-test. Statistical analysis was performed in GraphPad Prism 9, error bars show standard error of the mean (SEM).

## Acknowledgements

We gratefully acknowledge the expert technical assistance of Mette Flinck. Ester E. Sørensen, Kristian Lindholm, and Dan P. Christensen contributed with early pilot experiments and discussions. The primary cortical mouse neurons were kindly prepared and provided by the group of Blanca I. Aldana. The N-terminal epitope NBCn1 antibody was a kind gift from Jeppe Prætorius, Aarhus University.

## Author contributions

Conceptualization, MS, MTB and SFP; Methodology, MS, ELP, MTB, DC. Investigation, MS, MTB, ELP, FBC, MP, IAX, TL, LP; Writing – Original Draft, MS and SFP; Writing – Review and Editing, MS and SFP, with inputs from all authors; Funding Acquisition, SFP; Resources, MS, MTB, SFP; Supervision, MS, SFP. All authors have seen and approved the last version of the manuscript.

## Data availability

All data are available from the authors upon reasonable request.

